# Developing an accessible judgement bias task for non-human primates

**DOI:** 10.64898/2026.09.28.754938

**Authors:** Aurore Serda, Ruta Vaicekauskaite, Charlotte Canteloup, Hélène Meunier

**Author notes:** Corresponding Author: Aurore Serda^a^, 12 rue Goethe, Strasbourg, 67000, France. Authors contributed equally to this work (Charlotte Canteloup, Hélène Meunier).

## Abstract

Graphical abstract

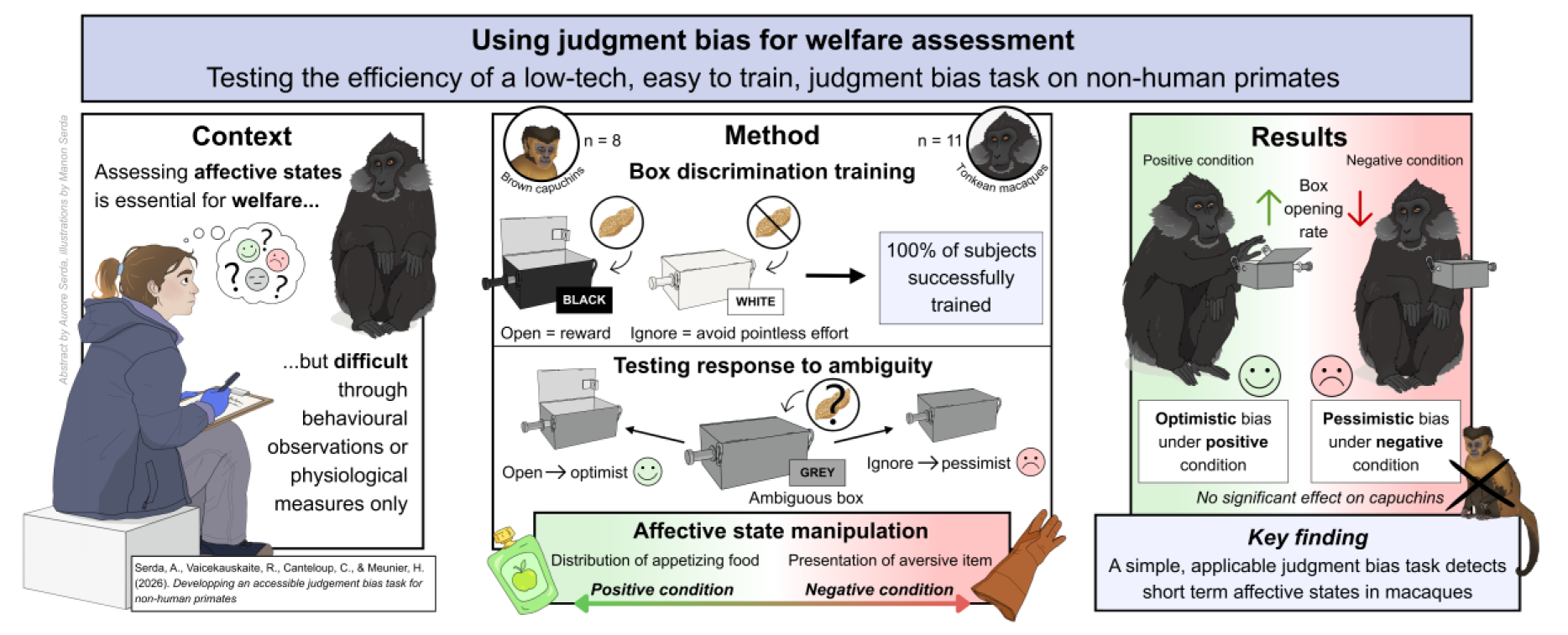

Objective and efficient methods for investigating animal’s mental states are important for advancing animal welfare research. Behavioural observations and physiological measures provide valuable information but may not fully capture affective states. Judgment Bias Tasks (JBTs) offer a promising method to infer affective states from animals’ responses to ambiguous stimuli, as distressed individuals show more pessimistic tendencies. However, the application of JBTs outside research settings may be limited due to time-consuming training and methodological or infrastructural requirements. With the present study, we aimed to test the feasibility of a simple and accessible JBT for non-human primates and to investigate its sensitivity to affective states. Eight brown capuchins (*Sapajus apella*) and eleven Tonkean macaques (*Macaca tonkeana*) were tested under positive and negative affective conditions. Individuals were trained to discriminate between never rewarded conditioned stimuli and always rewarded ones. They were then presented with ambiguous probes they had never seen before to analyse their responses to ambiguity. The discrimination training required, on average, about 12 sessions lasting a few minutes each to successfully train every individual animal. Both species significantly discriminated between conditioned stimuli and ambiguous probes. Additionally, macaques showed a greater tendency to interact with the never rewarded condition stimuli and the probes under positive conditions during early sessions. In capuchins, however, the affective conditions presented had no significant effect, although the descriptive analysis tended to show that the latency to investigate the probes was greater in the negative condition. This study also highlights key methodological considerations, such as a decrease in ambiguity across sessions. Overall, the present results suggest that, with a few improvements, this method could be effectively implemented to assess affective states of primates under human care, supporting this tool as a valuable candidate for welfare assessment.

**Highlights:**

- Low-tech, easy to implement, fast to train judgment bias task
- All 19 individuals were successfully trained in less than 12 sessions on average
- Macaques, but not capuchins, showed optimism under a positive affective state

## Introduction

The welfare of non-human animals (hereafter referred to as ‘animals’) has been the subject of growing attention over the past decades (Fraser et al., 1997; Marinou and Dontas, 2023; Sinclair et al., 2022). Advances in our knowledge and understanding of animal’s subjective experiences have highlighted the importance of carefully assessing animal welfare wherever animals are housed (Reimert et al., 2023). This can be challenging, as housing conditions, availability of the staff, time constraints, and other practical considerations can vary considerably across institutions (e.g. laboratories, zoos, shelters) depending on their resources, goals or infrastructures. Most modern assessments of animal welfare are based on the Five Domains Model (Mellor et al., 2020), a framework that takes into account four physical/functional domains: nutrition, environment, physical health and behavioural interactions - all of which are factors contributing to a fifth domain: the mental state. The mental state domain, which refers to the animal’s overall subjective internal experience, is difficult to assess. When addressing this domain, research generally focuses on *affective states,* as defined by Mendl and Paul (2020), as an internal state, with a positive or negative valence, that is not necessarily experienced consciously. Behavioural (Altmann, 1974) and physiological (DeSana et al., 2024; Wascher, 2021) measurements are commonly used to study this mental domain but they may prove insufficient to fully characterize an animal’s affective state. Inter-observer reliability issues (Burghardt et al., 2012), understudied species (Rosenthal et al., 2017), underrepresented behaviours (Wark et al., 2021), training requirements (Bloomsmith et al., 2015) or environmental conditions (Heintz et al., 2019; Ross et al., 2021) may also limit their use and accuracy.

Emotions have been reported to impact animal’s cognitive abilities (Blanchette et al., 2017; Boissy and Lee, 2014), notably how individuals evaluate ambiguous situations (Roelofs et al., 2016). This is the famous glass ‘half full or half empty’ analogy: depending on how one perceives this metaphorical glass, one is considered more pessimistic or optimistic. This is known as judgment bias (Lagisz et al., 2020). In the context of this bias, pessimism is defined as a relatively lower expectation of reward or higher expectation of punishment (optimism, the reverse; Bateson, 2016). Such operationalized measures of pessimistic and optimistic tendencies have been used to infer an individual’s affective state (Lagisz et al., 2020) through ‘Judgment Bias Tasks’ (JBTs). In these tasks, animals are typically trained to discriminate a rewarded (CS+) from a punished or non-rewarded (CS-) stimulus. They are then tested with untrained ambiguous probes (P). Judgment bias theory predicts that positively valenced affective states will lead to more optimistic responses to these probes (responses similar to CS+), while negatively valenced states will lead to more pessimistic responses (similar to CS-responses). JBTs have been used across a wide range of species, from rodents to birds, from farm animals to primates (Bethell, 2015; Mendl et al., 2009; Roelofs et al., 2016). Animals living in enriched environments tend to make more optimistic judgements (Bethell and Koyama, 2015; Matheson et al., 2008), while those displaying stereotyped behaviours, receiving higher rates of aggression or being tested after veterinary interventions tend to be more pessimistic (Bethell et al., 2012; Brilot et al., 2010; Gordon and Rogers, 2015; Pomerantz et al., 2012). Overall, JBTs have garnered considerable interest as a tool for assessing animal welfare, serving as an indirect measure of affective state (Bethell, 2015; Bethell and Pfefferle, 2023; Clegg, 2018).

Although JBT studies have yielded promising results, some concerns remain about their practical application (Lenkei et al., 2026). Some tasks require a large number of training sessions, sometimes exceeding 20 (e.g. Ash and Buchanan-Smith, 2016; Gordon and Rogers, 2015; Starling et al., 2014; Winkler et al., 2025) or 40 sessions (e.g Bethell et al., 2012; Verbeek et al., 2019); others are characterized by a significant proportion of individuals who do not meet the training criteria (e.g. Bethell and Koyama, 2015; Hintze et al., 2017; Perea-Rodriguez et al., 2024; Starling et al., 2014; Verbeek et al., 2019; Wichman et al., 2012). It may therefore be difficult to replicate these methodologies, as animal-care staff may lack the necessary time and resources, and standardized welfare assessments generally require that each individual be evaluated. Some of the devices used in JBTs may also not be available in all settings (touchscreens, for example; Bethell et al., 2012). Furthermore, studies sometimes lack clear empirical methodological guidelines (optimal number of sessions, probes, trials, and their order), despite evidence that repeated, unrewarded exposure to ambiguous probes can reduce their ambiguity as testing progresses (Brilot et al., 2010; Doyle et al., 2010; Wilson et al., 2023).

With this study, we sought to develop a JBT that is accessible, effective, and quick-to-train. We conducted this study on two species of non-human primates (hereafter ‘primates’): brown capuchin (*Sapajus apella*), a species previously successfully tested in JBT research (Pomerantz et al., 2012; Schino et al., 2016), and Tonkean macaque (*Macaca tonkeana*), never tested before. Our criteria were that this task should be accessible to the vast majority, if not all, of the individuals tested, as well as to different species. We also ensured that the task was technically undemanding, so that it could be implemented under varying constraints. We used only two probes, limiting the amount of material and the number of trials needed. We conducted ten sessions for each probe to help identify an optimal number of presentations before the probes lost their ambiguity. We also evaluated two types of response variables: binary outcomes (relatively easy to record and analyse) and response latencies (requiring more detailed data collection and analysis). To assess the task’s sensitivity to affective state, we tested it under two induced affective conditions (positive or negative). We hypothesized that both species would discriminate the cues based on their position on the CS-to CS+ continuum, with interaction increasing as cue valence increased (CS-< P-< P+ < CS+). If this paradigm is sensitive to affective state, we predicted that individuals would interact more quickly and frequently with P- and P+ under positive than negative conditions. Furthermore, we expected an effect of time, as animals might learn, over the course of the sessions, that the ambiguous probes were not reinforced.

## Materials & Methods

### Ethical statement

All procedures were non-invasive. The animals’ participation was always voluntary; individuals were never restrained or forced to interact with the stimuli. No food or water restriction was imposed. This research was approved by the Ethics committee of the Simian Laboratory Europe (SILABE) - University of Strasbourg (SBEA), which is authorized to house non-human primates (registration n° B6732636). The research also complied with the EU Directive 2010/63/EU for animal experiments.

## 1. Animals and Housing

The study took place between May and July, and November and December of 2025, at the SILABE - University of Strasbourg. Twenty individuals were initially recruited but one was excluded before testing as his willingness to participate in training sessions was highly unreliable (see section Results 1. Training). Our study therefore included 19 individuals: 11 Tonkean macaques (mean age = 11.8 years, range 3-26; 10 males, one female) and eight brown capuchins (mean age = 12.3 years, range 8-23; six males, two females; Supplementary Table S1). The animals were living in seven different groups (four macaque groups, ranging from two to 20 individuals; three capuchin groups, ranging from two to three individuals). Some groups were family groups, including males, females, adults, and juveniles, while others consisted exclusively of adult males or adult females. Tonkean macaque groups were housed either in large naturalistic forested outdoor enclosures or in smaller outdoor enclosures on concrete floor. Brown capuchins were housed in small outdoor enclosures with concrete or dirt floor (see Supplementary Table S1 for details). All groups had permanent access to indoor facilities, *ad libitum* water, a daily ration of pellets, and fresh fruits and vegetables once a week. Tests were conducted in the enclosure or in a dedicated adjacent area, either alone or in the presence of conspecifics, handled by an experimenter to prevent any interference.

## 2. Experimental design

### 2.1. Go/No-go paradigm

We opted for a ‘Go/No-go’ paradigm in which animals were presented with a stimulus (‘CS+’, ‘CS-’ or ambiguous probes ‘P’) and had to choose either to interact or to ignore it. ‘Go/Go’ paradigms (in which the animal always interacts but must choose between two different cues) were considered, but their more demanding training requirements made them less suitable. ‘Go/No-go’ tasks, however, still pose training challenges, such as the fact that some individuals always or never interact with the stimuli, regardless of the type of signal (Bateson and Nettle, 2015). We hypothesized, however, that some of these behaviours might reflect motivation-related effects rather than lack of understanding. Thus, the failures observed during training could be partly explained by the absence of a cost associated with interacting with the stimuli. Based on this hypothesis, we sought to design a task requiring a minimal effort threshold while ensuring rapid accessibility. We predicted that this would encourage individuals to refrain from systematically interacting with the CS-.

### 2.2. Apparatus

The stimuli used were closed boxes that contain or not a food reward (a peanut for all animals except two who received raisins). The boxes were made of metal, rectangular (11 × 7 × 7.5 cm), and equipped with a side handle that the individuals had to pull to the side before lifting the lid to open the box (see Figure 1). The boxes could be easily attached to and removed from the outside of different enclosures using carabiners, allowing the animals to interact by reaching their hands through the mesh. Four types of boxes were used. A black box that always contained a food reward served as the positive conditioned stimulus (CS+). A white box, which was always empty, served as the negative conditioned stimulus (CS-). Two grey boxes, which were always empty, served as the ambiguous probes (P): a light grey box was the ‘probe near the negative’ (P-), and a dark grey box was the ‘probe near the positive’ (P+). The colours of the grey boxes were calibrated to be approximately equidistant from the black and white boxes (light grey at perceptual position 0.33 on a scale ranging from 0 (white) to 1 (black); dark grey at position 0.66; see ‘Supplementary Methods S1’ for the calculation method).

**Figure 1:**
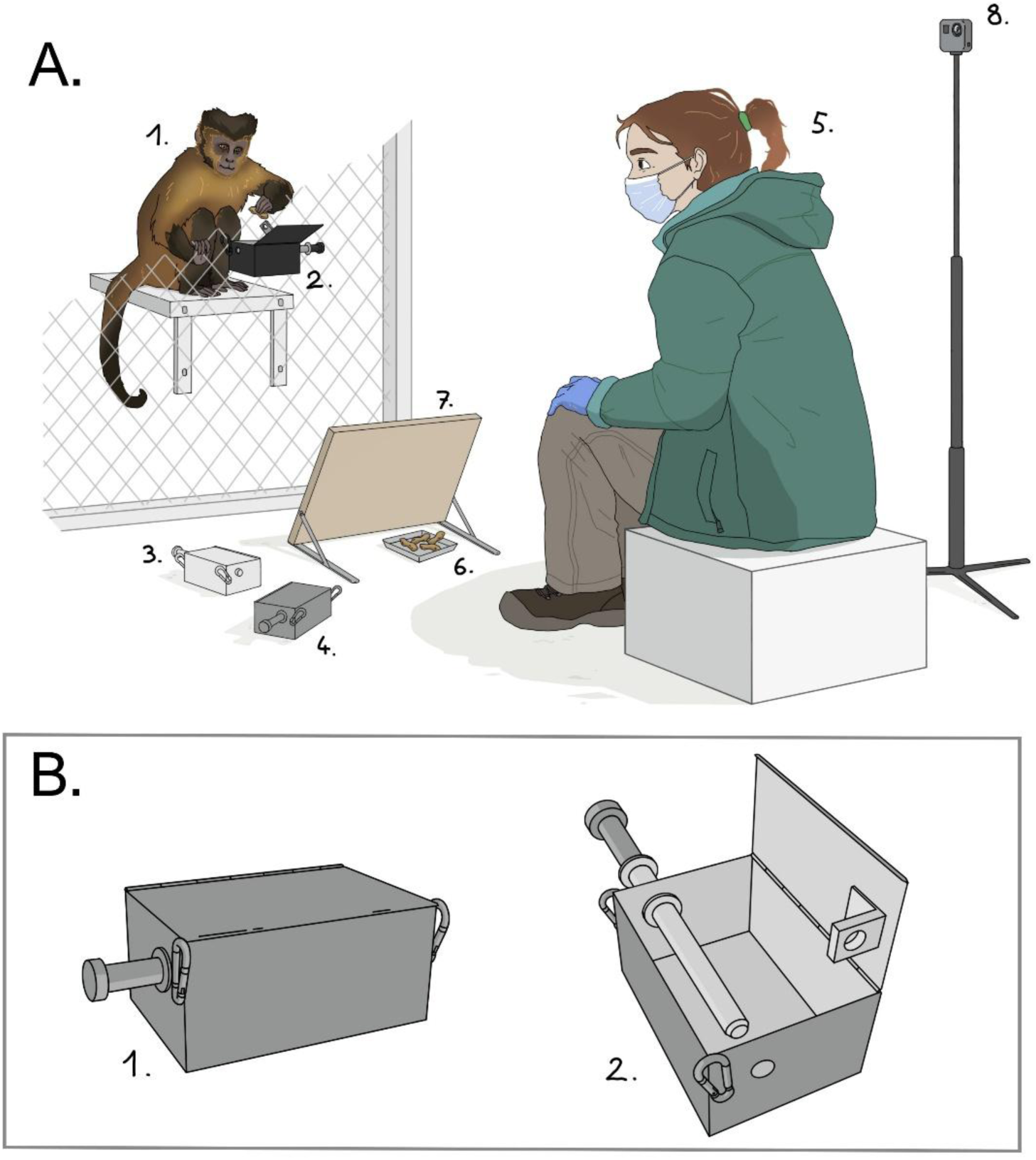
(A) Schematic representation of a test session: the subject (1) opens the CS+ (2) and grabs the reward. The other two boxes, CS-(3) and P (4), are ready to be presented by the experimenter (5) in subsequent trials. Rewards (6) are hidden behind a screen (7) so that the experimenter can refill the CS+ after a trial. The session is filmed by a Gopro camera (8). (B) Schematic representation of the apparatus closed, from the subject’s facing side (1) and opened (2). The metallic rod running across the box had to be pulled to the side before the lid could be opened. Illustration by Manon Serda.

## 3. Training phase

### 3.1. Box manipulation training

The individuals were first trained on how the box worked by learning to pull the handle and lift the lid of the black box that was consistently rewarded (CS+). Each training session consisted of six trials, and only one session was conducted per day. The box was presented to the individuals, who were free to interact with it. If an individual did not spontaneously pull the handle after a few minutes, the experimenter assisted them until they were able to perform the task independently. An individual was considered to have successfully completed this phase when they consistently opened the black box without assistance in 100% of trials during two consecutive sessions.

### 3.2. Cue discrimination training

Individuals who successfully completed the first training phase were then trained to associate the colour of the box (black or white) with the presence or absence of a reward. Each session consisted of 12 trials: six presentations of the black box (CS+) and six presentations of the white box (CS-), in a pseudo-randomized order (no more than two consecutive presentations of the same colour). During each trial, the box was accessible for a maximum duration of five seconds. This duration was based on the typical time observed for an individual to decide whether to interact with the box or ignore it. If the individual did not attempt to open the box during this time, it was removed and the trial was recorded as ‘unopened’. If the individual initiated an interaction with the box within the allotted time, they had unlimited time to complete the task, provided they remained actively engaged. If they disengaged for more than five consecutive seconds, the box was removed, and the trial was recorded as unopened. Once a box was opened, it was immediately removed after the individual finished exploring it. The interval between trials depended on the outcome of the previous trial, with white boxes resulting in a time penalty (20 seconds) if opened (see Supplementary Figure S1 for a schematic representation of the decision-making process). Individuals successfully completed this phase when they opened at least five of the six black boxes and no more than two of the six white boxes per session, over the course of two consecutive sessions. Importantly, individuals were never exposed to the grey boxes during training, which ensured their ambiguity at the start of the test phase.

Two individuals deviated from this criterion because they completed additional sessions after they had already met it. This was because methodology was more demanding in early training but was revised to avoid unnecessary overtraining of individuals. However, those individuals were included in the main analysis, and the results proved robust even to their exclusion (Supplementary Table S2). Another individual underwent a retest to assess the durability of the training, as the criterion had been validated three weeks before the start of the tests. The individual validated two of the three retest sessions. The failure during the second session followed an episode of collective aggression and coincided with abnormal reactivity; the criterion was therefore validated based on the other two sessions.

## 4. Testing phase

### 4.1. Phases

The tests were conducted in two successive phases, each consisting of 10 sessions. During the first phase, known as the ‘light phase’ (phase L), the light-grey box (P-) was tested on the entire cohort (n = 19). The second phase, known as the ‘dark phase’ (phase D), was conducted following phase L and involved testing the dark-grey box (P+) on a subgroup of the cohort (n = 13). This phase could not involve all individuals of phase L due both to time constraints and because some animals had left the facility. Phase D was primarily designed as a control to assess whether individuals perceived ambiguity along a continuum ranging from CS-to CS+ and whether the probes were not simply perceived as a random colour. The two probes were therefore tested during separate phases rather than during the same sessions. This design allowed phase L to serve as the main test to determine whether it was possible to detect a judgment bias using a single ambiguous probe.

### 4.2. Procedure

Each individual was tested in two different conditions (positive and negative; see below section 4.3), over the course of five sessions per condition, with a maximum of one session per condition per week. Each session consisted of nine trials: three presentations of the black box (CS+), three presentations of the white box (CS-), and three presentations of the grey box (P). Trial order was pseudo-randomized, with the same box never being presented in two consecutive trials. The sessions followed the same procedure as the training phase, with each box, including the grey box, being presented sequentially and for a maximum of five seconds. The grey box was never rewarded; if it was opened, it resulted in a 10 second delay before the next trial (Supplementary Figure S1). All sessions were video recorded using a GoPro HERO10 (GoPro Inc., San Mateo, CA, USA).

### 4.3. Conditions

The testing phase was conducted under two conditions: a positive condition and a negative condition. In both conditions, the animal voluntarily entered the testing area. On the one hand, in the positive condition, the experimenter offered the individual approximately 15g (for capuchins) to 20g (for macaques) of applesauce (a highly appetizing food for all individuals) and once the individual had eaten and was attentive and ready, the test session began. On the other hand, in the negative condition, the individual was presented with an object designed to cause mild discomfort. The object varied depending on individual reactions (capture glove, partially concealed capture net, plastic bag). The session began immediately after the object was presented, as soon as the individual was attentive and ready.

## 5. Data analysis

Behavioural responses were coded from video recordings using the BORIS software (v. 8.27.10, Friard and Gamba, 2016). For each trial, the latency to the first opening attempt was recorded (in seconds, ranging from zero to five seconds). This corresponded to the time elapsed between the presentation of the box and the first attempt to open it. Trials in which no interaction occurred within the five-second time frame were assigned a latency value of five seconds, which allowed all trials to be included in the latency analyses. Opening rates were also recorded, as a binary score: 1 = box opened; 0 = box not opened. A random subset of 20% of the videos was selected for double coding. Inter-rater reliability was assessed using Cohen’s Kappa coefficient (all Kappa > 0.80). Coders could not be blind to experimental conditions.

## 6. Statistical analysis

All analyses were conducted in R v4.3.2 (R Core Team, 2025). Table 1 provides an overview of all models. The probability of opening the box (binary variable: opened/not opened) was modelled using generalised linear mixed models (GLMM; glmmTMB package; Brooks et al., 2017) with a binomial error distribution and logit link function. The latency to open a box (in seconds, right-censored at five seconds for trials without interaction) was modelled using mixed-effects Cox proportional-hazards models (coxme package, Therneau, 2009). A hierarchical random-effect structure (session number nested within subject) was specified to reflect the repeated-measures design.

**Table 1:** Overview of the statistical models. Model names follow the convention MODEL[N]_[Species], where N indicates the research question (1 = cue discrimination; 2 = affective state effect; 3 = within-session repetition), and Species is M (Macaca tonkeana) or S (Sapajus apella). For question 2, an additional suffix (_L or _D) indicates the testing phase (Light or Dark). All the models included random effects (1|Subject/sessionNBR). For Cox models, N obs is reported as events (interaction with the box) / total trials.

| Question | Model name | Distribution | Response variable | Phase | Fixed predictors | Data subset | N obs / N subjects<br>*M. tonkeana* | N obs / N subjects<br>*S. apella* |
| --- | --- | --- | --- | --- | --- | --- | --- | --- |
| **1. Cue discrimination:** do subjects distinguish between cue types? | GLMM1_M/<br>GLMM1_S | Binomial | Box opening | Both | Cue (categorical) + session number + first trial | All cues | 1346 / 11 | 1418 / 8 |
|  | Cox1_M/<br>Cox1_S | Cox proportional hazards | Latency to interact with the box |  | Cue (categorical) × session number + first trial | All cues | 736 events / 1346 / 11 | 878 events / 1418 / 8 |
| **2. Affective state effect:** does condition affect responses to the different cues? | GLMM2_M_L/<br>GLMM2_S_L | Binomial | Box opening | Phase L | *M. tonkeana:*<br>Condition × cue (continuous) × session number + first trial<br>*S. apella:*<br>Condition + cue (continuous) + session number + first trial | CS-, P-, CS+ | 950 / 11 | 716 / 8 |
|  | GLMM2_M_D/<br>GLMM2_S_D |  |  | Phase D | Condition + cue (categorical) + session number + first trial | CS-, P+, CS+ | 396 / 5 | 702 / 8 |
|  | Cox2_M_L/<br>Cox2_S_L | Cox proportional hazards | Latency to interact with the box | Phase L | Condition + cue (categorical) × session number + first trial | CS-, P-, CS+ | 512 events / 950 / 11 | 469 events / 716 / 8 |
|  | Cox2_M_D/<br>Cox2_S_D |  |  | Phase D | Condition + cue (categorical) × session number + first trial | CS-, P+, CS+ | 224 events / 396 / 5 | 409 events / 702 / 8 |
| **3. Within-session repetition:** does responding decline across presentations of the same cue? | GLMM3_M /<br>GLMM3_S | Binomial | Box opening | Both | Cue (categorical) × presentation number + session number + first trial | All cues | 1346 / 11 | 1418 / 8 |

The cue type was coded either as a four-level categorical predictor (CS-, P-, P+, CS+) or as a continuous variable reflecting the colorimetric position measured along an axis ranging from CS-to CS+ (0, 0.3, 0.7, 1, Supplementary Methods S1), in accordance with the recommendations of Gygax (2014). For each GLMM, the choice between a continuous or a categorical variable was based on a comparison using the Bayesian Information Criterion (BIC). Categorical coding was used for all Cox models to ensure consistency with Kaplan-Meier plots. This aligned with BIC results for Cox models of phase D and was a moderate trade-off for phase L models which showed slightly higher BIC for categorical models. Since no a priori hypothesis was made regarding differences between species, the models were therefore fitted separately for each species. Since the ambiguous P- and P+ probes were tested during separate experimental phases and were never presented during the same session for a given subject, they are structurally confounded with the test phase. This confounding factor was avoided for the affective-state models by fitting separate models for each phase (Table 1). However, it was retained for the discrimination models, where the explicit objective was to make a direct comparison between P-vs P+, which is recognized as a limitation in this context.

A control covariate related to the trial’s position within the session (first trial vs. others) was included in all models, based on preliminary analysis showing a consistently higher response rate during the first trials. To support practical recommendations regarding the protocol, an additional GLMM was used to examine whether the response rate decreased across successive presentations of the same cue within a single session. The significance of fixed effect was assessed using Type II Wald χ^²^ tests (car package; Fox and Weisberg, 2019). Non-significant interaction terms (p > 0.05) were removed (Gygax, 2014) to interpret simple effects. Estimated marginal means and pairwise comparisons (adjusted according to Tukey or Holm correction) were obtained using the emmeans function (Lenth, 2025) and trends were obtained using the emtrends function. When interactions were significant, contrasts were extracted at fixed points along the relevant predictors rather than interpreted based on the averaged main effects.

The fit of binomial model was assessed using simulated residuals (DHARMa package; Hartig, 2024). For survival models, the proportional-hazards assumption was assessed using the Grambsch-Therneau test (cox.zph) and Schoenfeld residuals. In accordance with the recommendations of Therneau & Grambsch (2000), significant test statistics were not considered to invalidate the models when graphical inspections revealed no problematic patterns and the sample size was large. A sensitivity analysis limited to individuals tested during both experimental phases was conducted for macaques (all capuchins completed both phases) and showed overall consistent results in terms of direction and significance (Supplementary Table S2). To assess the feasibility of individual-level interpretation, the direction and magnitude of the condition’s effect on the opening of boxes and on the latency to interact with P-was examined descriptively for each subject.

### Data availability statement

All data and code used in this study are available at: https://doi.org/10.5281/zenodo.22938479

## Results

Nineteen individuals (11 Tonkean macaques and eight brown capuchins) were trained and tested on a Go/No-go judgement bias task under positive and negative induced affective conditions. Results are presented in three sections: training phase, testing phase (including cue discrimination and the effect of affective condition) and practical considerations regarding session and trial number.

## 1. Training phase

On average, all the 19 individuals required 4.95 ± 2.34 sessions (range: 2-10) amounting to 29.68 ± 14.07 trials (range: 18-60) to meet the criterion for mastery of box manipulation.

Regarding box discrimination, all the 19 individuals required an average of 11.89 ± 3.62 sessions (range: 6-18) or 142.74 ± 43.43 trials (range: 72-216) to meet the criterion.

## 2. Testing phase

A total of 2764 trials of the judgment bias task were recorded (96.0% of the expected 2880 total across n = 19 individuals). Individuals completed on average 9.79 ± 0.12 sessions in phase L (n = 19) and 9.38 ± 0.27 sessions in phase D (n = 13).

### 2.1. Cue discrimination

#### Proportion of box-opening

The type of cue had a significant effect on the probability of opening the box in both macaques (GLMM1_M: Type II Wald χ^²^ = 242.45, p < 0.001) and capuchins (GLMM1_S: χ^²^ = 226.27, p < 0.001). Whether it was the first-trial or not also had an effect on box opening in both species (GLMM1_M: χ^²^ = 64.65, p < 0.001; GLMM1_S: χ^²^ = 62.28, p < 0.001) with higher opening probabilities during the first trial of a session (GLMM1_M: β = 2.64, SE = 0.33, p < 0.001; GLMM1_S: β = 2.60, SE = 0.33, p < 0.001). The session number did not have a significant effect in macaques (GLMM1_M: χ^²^ = 1.44, p = 0.230) but it did in capuchins (GLMM1_S: χ^²^ = 11.44, p < 0.001), in which the probability of opening the box declined over the course of the sessions (slope = −0.30, SE = 0.09, p < 0.001).

The estimated mean probabilities (at first_trial = 0) followed the expected hierarchy in both species, with CS-boxes being opened less than P-boxes, P-boxes less than P+ boxes and P+ boxes less than CS+ boxes (Figure 2a; Supplementary Table S3). In macaques, all pairwise contrasts between cues were significant (all p < 0.01; Table 2); in capuchins, all contrasts involving CS- or CS+ were significant (all p < 0.001), but P-did not differ significantly from P+ (p = 0.735, see Table 2).

**Figure 2:**
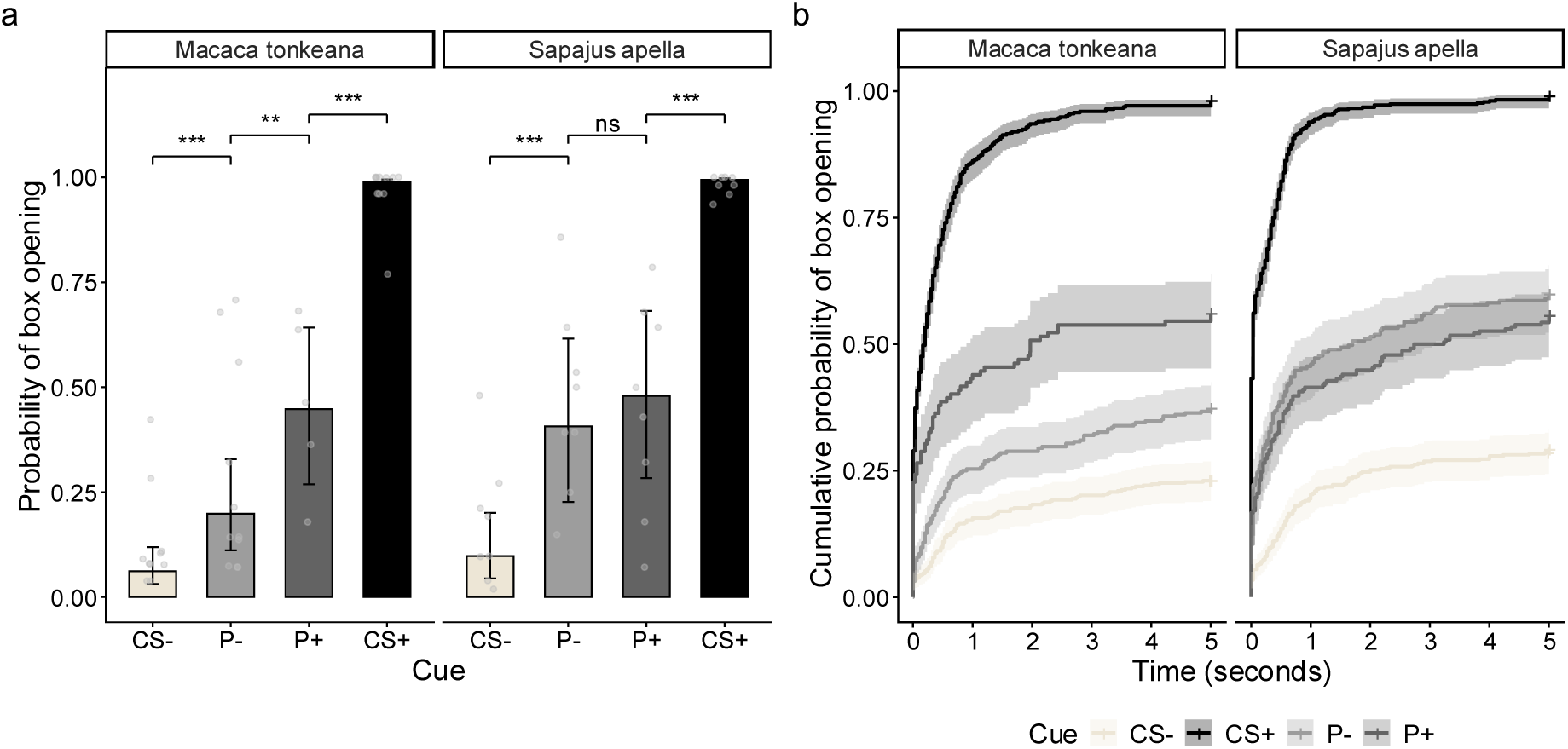
Cue discrimination by species. (a) Model-estimated probability of box-opening by cue (GLMM1_M, GLMM1_S), with individual subject means (points) and pairwise significance indicators (Tukey-adjusted; ** p < 0.01, *** p < 0.001). (b) Cumulative probability of box-opening over time (0-5 s), by cue type, based on Kaplan-Meier estimation; shaded areas indicate 95% confidence intervals. CS-= negative conditioned stimulus; P-= probe closer to CS-; P+ = probe closer to CS+; CS+ = positive conditioned stimulus.

**Table 2:**
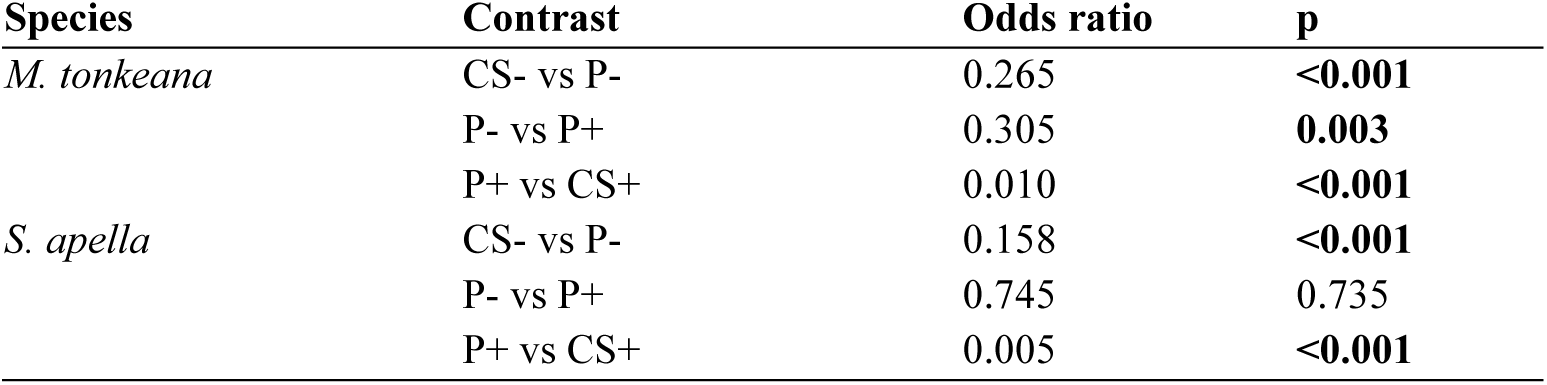
Post-hoc pairwise comparisons of estimated marginal means for cues by species, along the ambiguity gradient (GLMM1_M, GLMM1_S; Tukey-adjusted). Remaining pairwise contrasts (CS-/P+, CS-/CS+, P-/CS+) were all significant at p < 0.001 in both species.

#### Latency to open the box

For both species, the effect of the interaction between cue and session (Cox1_M; χ^²^_₃_ = 23.12, p < 0.001; Cox1_S; χ^²^_₃_ = 28.61, p < 0.001) and of the first trial on the latency to open the box were significant, with faster opening on the first trial of a session (Cox1_M: β = 0.45, p < 0.001; Cox1_S: β = 0.51, p < 0.001). Due to the interaction between cue and session, the ambiguity hierarchy among cues was assessed using pairwise contrasts extracted from each session rather than from the averaged slopes (Table 3).

**Table 3:** Pairwise contrasts of latency to open the box along the ambiguity gradient (consecutive comparisons), by session and species, from the mixed-effects Cox model (Cox1_M, Cox1_S; Holm-adjusted). Each cell reports hazard ratio / SE / p-value. Ratios above 1 indicate faster opening for the first-listed cue. Remaining contrasts (CS- vs CS+, CS- vs P+, P- vs CS+) were significant at p < 0.001 at every session in both species and are available in Supplementary Table S4.

| Species | Contrast | Session 1 | Session 2 | Session 3 | Session 4 | Session 5 |
| --- | --- | --- | --- | --- | --- | --- |
| *M. tonkeana* | P- / CS- | 1.62 / 0.35 / 0.053 | 1.91 / 0.29 / **<0.001** | 2.25 / 0.33 / **<0.001** | 2.65 / 0.53 / **<0.001** | 3.12 / 0.88 / **<0.001** |
|  | P+ / P- | 1.67 / 0.47 / 0.067 | 1.73 / 0.33 / **0.005** | 1.78 / 0.31 / **0.001** | 1.84 / 0.43 / **0.010** | 1.90 / 0.65 / 0.059 |
|  | CS+ / P+ | 3.39 / 0.83 / **<0.001** | 4.05 / 0.69 / **<0.001** | 4.84 / 0.74 / **<0.001** | 5.79 / 1.21 / **<0.001** | 6.92 / 2.07 / **<0.001** |
| *S. apella* | P- / CS- | 3.23 / 0.64 / **<0.001** | 3.14 / 0.45 / **<0.001** | 3.06 / 0.38 / **<0.001** | 2.97 / 0.47 / **<0.001** | 2.89 / 0.64 / **<0.001** |
|  | P+ / P- | 1.27 / 0.27 / 0.254 | 1.06 / 0.16 / 0.698 | 0.88 / 0.12 / 0.350 | 0.74 / 0.13 / 0.079 | 0.61 / 0.15 / **0.047** |
|  | CS+ / P+ | 2.47 / 0.42 / **<0.001** | 3.62 / 0.45 / **<0.001** | 5.29 / 0.60 / **<0.001** | 7.75 / 1.16 / **<0.001** | 11.34 / 2.38 / **<0.001** |

During all sessions, macaques opened CS+ boxes significantly faster than P+ boxes (Table 3). Furthermore, during sessions 2 through 4, macaques opened P+ boxes significantly faster than P-boxes, and during sessions 2 through 5 P-boxes faster than CS-boxes. In capuchins, similarly, CS+ boxes were opened significantly faster than P+ boxes and P-boxes were opened faster than CS-boxes. However, no difference in latency to open the box was significant between P- and P+ at the exception of the fifth session where P+ boxes were opened more slowly than P-boxes.

The Kaplan-Meier curves confirmed this stable hierarchy in both species, with the cumulative probability of opening the P- and P+ boxes across time completely overlapped in capuchins (Figure 2b).

### 2.2. Effect of affective states on boxes’ opening

#### Proportion of box-opening

A significant effect of the condition (positive; negative) on the probability of opening the box was found only in macaques during phase L (P-): the three-way interaction of condition×cue(continuous)×session_number was significant (GLMM2_M_L: χ^²^_₁_= 5.30, p = 0.021). Comparison between conditions was examined for each cue position (0 = CS-, 0.3 = P-, 1 = CS+) and session (Holm-adjusted) and showed that the effect of condition was strongest and most consistent in the first sessions. CS- and P-boxes were opened significantly more under the positive than negative condition during sessions 1, 2 (both p < 0.01), and 3 for P-boxes (p = 0.035), with no significant effect of condition for CS+ in any session (all p > 0.20; see Table 4 and Figure 3a). A significantly greater decrease in the probability to open the CS-boxes was found across sessions under the positive condition than the negative one (negative condition: slope = 0.20, SE = 0.20, p = 0.315; positive condition: slope = −0.53, SE = 0.19, p = 0.005; slope difference = 0.73, p = 0.008), with a similar but non-significant trend for P-boxes (negative: slope = 0.11, SE = 0.16, p = 0.482; positive: slope = −0.25, SE = 0.15, p = 0.106; slope difference = 0.36, p = 0.106). No such difference was observed for CS+ boxes (slope difference = p = 0.240).

**Figure 3:**
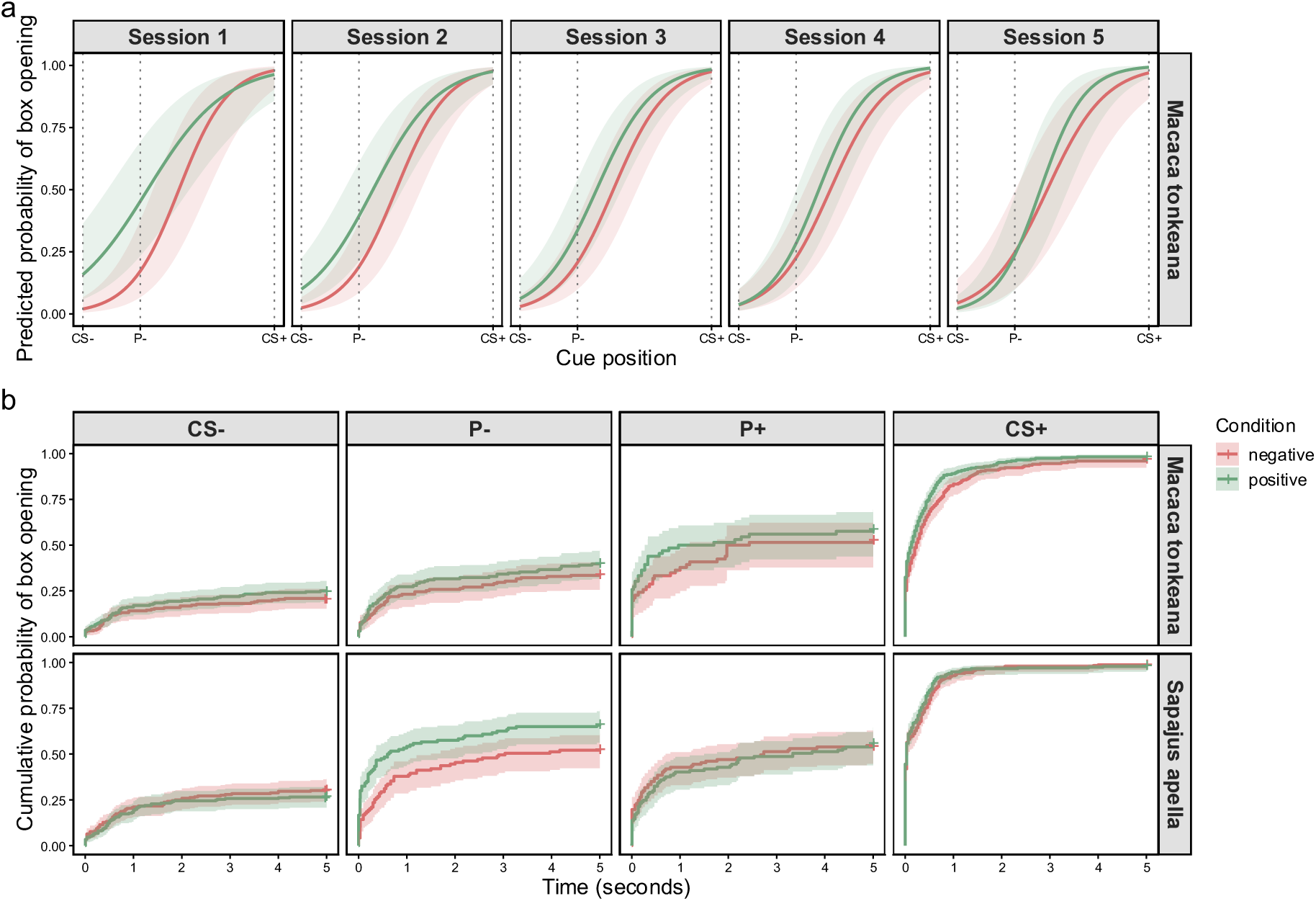
Effect of affective state (negative or positive condition). (a) Model-predicted probability of box-opening in macaques (phase L; GLMM2_M_L) as a function of cue position (0 = CS-, 0.3 = P-, 1 = CS+) and condition, across sessions 1-5; shaded areas indicate 95% confidence intervals, dotted vertical lines indicate observed cue positions. Only macaques are shown, as this was the only species×phase combination with a significant condition×cue×session interaction; condition had no significant effect in the remaining three models (macaques phase D, capuchins phases L and D). (b) Cumulative probability of box-opening over time (0-5 s), by condition, species, and cue, based on Kaplan-Meier estimation; shaded areas indicate 95% confidence intervals.

**Table 4:** Condition comparison (positive vs. negative) at fixed cue positions and sessions, macaques phase L, from the binomial GLMM (GLMM2_M_L; condition × cue_position × session_num interaction; Holm-adjusted). Each cell reports odds ratio / SE / p-values. Ratios above 1 indicate higher opening probability under positive condition.

| Cue position | Session 1 | Session 2 | Session 3 | Session 4 | Session 5 |
| --- | --- | --- | --- | --- | --- |
| CS- (0) | 9.33 / 6.14 / **<0.001** | 4.48 / 2.10 / **0.001** | 2.15 / 0.87 / 0.058 | 1.03 / 0.52 / 0.951 | 0.50 / 0.35 / 0.324 |
| P- (0.3) | 3.98 / 2.10 / **0.009** | 2.78 / 1.03 / **0.006** | 1.94 / 0.61 / **0.035** | 1.36 / 0.54 / 0.444 | 0.95 / 0.53 / 0.921 |
| CS+ (1) | 0.55 / 0.53 / 0.536 | 0.92 / 0.63 / 0.897 | 1.53 / 0.94 / 0.490 | 2.56 / 2.09 / 0.250 | 4.28 / 4.94 / 0.209 |

In the other three models (macaques, phase D; capuchins, phase L; capuchins, phase D), no interaction involving the condition was significant, and the condition had no significant main effect on the boxes’ opening proportion (GLMM2_M_D: χ^²^_₁_ = 0.58, p = 0.444; GLMM2_S_L: χ^²^_₁_ = 0.15, p = 0.697; GLMM2_S_D: χ^²^_₁_ = 0.92, p = 0.337).

#### Latency to open the box

In all four models, as the three-way interaction ‘condition×cue×session’ was non-significant, it was removed from the models (Cox2_M_L: χ^²^_₂_ = 0.63, p = 0.730; Cox2_M_D: χ^²^_₂_ = 0.45, p = 0.799; Cox2_S_L: χ^²^_₂_ = 0.03, p = 0.987; Cox2_S_D: χ^²^_₂_ = 1.16, p = 0.559). In the simplified models, the condition had a significant main effect on the latency to open all boxes only in macaques during the phase L (Cox2_M_L: χ^²^_₁_ = 6.07, p = 0.014), with faster opening under the positive condition (negative vs. positive: ratio = 0.725, p = 0.014, averaged across cue and session). In capuchins, during phase L, this effect was a trend (χ^²^_₁_ = 2.89, p = 0.089). However, inspection of Kaplan-Meier curves (Figure 3b) revealed a consistent difference in the cumulative probability of opening the P-boxes across time between conditions, with boxes being opened faster under the positive condition. To determine whether this apparent pattern was driven by outliers, we examined the individual latencies for P-trials in greater details (Supplementary Table S5) and found that 7 of the 8 subjects had shorter latencies under the positive condition than the negative one (mean reduction of 33%, range: 7%-54%) with no individual exhibiting markedly higher latency variability than the others (SD range: 1.31-2.52 s). In phase D, the condition had no significant effect on the latency to open the boxes neither in macaques, (Cox2_M_D: χ^²^_₁_ = 0.16, p = 0.685) nor in capuchins (Cox2_S_D: χ^²^_₁_ = 0.03, p = 0.854).

## 3. Practical considerations

### 3.1. Within-session cue repetition

The probability of opening the boxes decreased not only across sessions but also within a session as the same cue was presented several times. In macaques, this decrease varied depending on the cue (GLMM3_M: χ^²^_₃_ = 10.20, p=0.017): it was significant for CS- and P-boxes’ opening rates (both p < 0.001, e.g. the opening of P-boxes decreased from 39% to 18% to 7% across presentations 1-3, see Supplementary Table S6), it was marginal for P+ boxes (p = 0.101), and nonsignificant for CS+ boxes (p = 0.837). In capuchins, the decrease did not differ by cue (GLMM3_S: χ^²^_₃_ = 7.44, p = 0.059) but was overall significant (GLMM3_S: χ^²^_₁_ = 15.62, p < 0.001).

### 3.2. Individual-level assessment

Due to experimental design constraints, i.e. low number of trials per individual to prevent probe familiarisation, the effects of condition at the individual level can be assessed only from a descriptive standpoint. The expected trend (higher P-boxes opening probability / shorter latency to open P-boxes in the positive condition) was observed in 7/11 (64%) macaques and 4/8 (50%) capuchins for opening probability, and in 8/11 (73%) macaques and 7/8 (88%) capuchins for latency (Supplementary Table S5).

## Discussion

The aim of this study was to design and to test the efficiency of a judgment bias task in the context of assessing primate welfare under human care. This study is one of the few in which all individuals successfully completed the training phase in a relatively small number of sessions. Both Tonkean macaques and brown capuchins effectively distinguished the ambiguous probes from the conditioned stimuli. However, only the macaques showed a clear discrimination between the two ambiguous probes and significantly opened more ambiguous boxes in the positive affective condition. Because the training and testing required only a relatively short investment in time, we believe this task is an efficient tool to assess short term affective states, at least in macaques. This study sheds light on a little-studied macaque species, the Tonkean macaque, and helps to expand the still very limited-but rapidly growing-body of primate research on judgment bias paradigms (see Machado et al., 2026 for a recapitulative table).

### Cue discrimination

All tested individuals successfully completed the discrimination training in less than twelve sessions on average. This is very encouraging as several studies on judgement bias report substantial numbers of subjects being unable to proceed to the testing phase because they failed the discrimination phase. For example, only half of subjects in mouse lemurs (*Microcebus murinus*; Perea-Rodriguez et al., 2024), none of three gorillas (*Gorilla gorilla gorilla*; McGuire et al., 2017; McGuire and Vonk, 2018), and only four of seven bonobos (Winkler et al., 2025) reached criterion after sometimes dozens of sessions or hundreds of trials. Compared to our task, some of the above-mentioned studies required more cognitively demanding conditional discrimination (McGuire and Vonk, 2018), or used only low-effort behavioural response, such as a simple approach or button press action (Perea-Rodriguez et al., 2024; Winkler et al., 2025). Our design combined a simple discrimination with a physically engaging response, what we believe could potentially explain this observed difference in training success. Although see Pomerantz et al. (2012) for a conditional discrimination and undemanding response with a 75% training success rate.

We also reported that individuals opened significantly less the negative conditioned stimulus than the ambiguous probes, which were also less opened than the positive conditioned stimulus. This confirms that subjects effectively distinguished the ambiguous probes from the conditioned stimuli. This expected gradient - the rate at which the boxes were opened following the negative-to-positive pattern of the cues - confirms that individuals perceived the probes as having intermediate valence, which further validates the design of our task.

Regarding the discrimination between the two probes, capuchins showed no differences in opening rate or latency, unlike the macaques. This decline in responses over repeated sessions is a common issue in judgement bias tasks using unrewarded probes (Brilot et al., 2010; Doyle et al., 2010; Perdue, 2017; Starling et al., 2014). We hypothesized that, since near-positive probes were introduced only during sessions 11-20, after the presentation of the near-negative probes (sessions 1-10), this decline should have continued and reduced the opening rates of the near-positive probe. Instead, near-positive probes were opened just as often as the near-negative ones, suggesting that they were interpreted more positively, offsetting the expected decline. The progression of the sessions did not affect discrimination in macaques, which is consistent with their slower initial learning, and could explain why they were not affected as quickly by the effects of the sessions and thus discriminated between all probes.

### Impact of affective states on responses to cues

In agreement with our hypothesis, we found that macaques opened more near negative probes (and opened all cues faster) under positive than negative affective condition in early sessions. This suggests that they are more optimistic under positively valenced affective state, and that those tendencies can be detected through simple binary responses. This is consistent with findings obtained in Japanese or rhesus macaques (*Macaca mulatta:* Bethell et al., 2012; *Macaca fuscata*: Iki and Adachi, 2023), and is encouraging for welfare assessment, as it suggests that even mild, short term affective states can be recognised through judgment bias tasks. Contrary to those studies which used touchscreens (Bethell et al., 2012; Iki and Adachi, 2023, 2025), we show that a very simple and low-tech paradigm can be very efficient. This is essential as zoos, sanctuaries and other facilities may have limited budget and resources to deploy high-tech testing techniques.

Interestingly, macaques also opened more negative conditioned stimuli in the positive condition, not only probes. This pattern is consistent with residual ambiguity of the negative conditioned stimulus (which was still occasionally opened, even after training). Indeed, if ambiguity was retained for this conditioned stimulus, it is expected that bias would impact it in a similar way it does the probe. Such effects have been previously reported in honeybees (*Apis mellifera carnica;* Bateson et al., 2011). Our results also show the positive conditioned stimulus to be unaffected by condition, ruling out the possibility of a general motivation effect (but see Strang and Muth, 2023 for a ‘peak-shift’ based alternative hypothesis).

In capuchins, although our descriptive analysis showed longer latencies for opening the near negative probes under negative condition, this effect was not significant in our models. Similarly, we report no significant effect of the induced affective condition on box opening rates. This lack of a condition-specific response may reflect a combination of task and species related factors, in addition to statistical power. We observed that capuchins opened the unrewarded boxes at a high rate, which limited the chances of detecting an effect. On the one hand, this could be explained by their more impulsive nature compared to macaques, who are better at tolerating waiting (Loyant et al., 2023; Pelé et al., 2011), for whom waiting the five seconds required before removing the box may have been relatively less costly. To avoid this, future studies could adjust the duration of the presentation of the boxes according to the species, rather than using a fixed duration (e.g. shorter presentation for capuchins to reduce the cost of ‘No-go’ responses). This could also explain why all but one individual showed the expected latency pattern, although nonsignificant, compared to only four individuals for opening rates: latency may better capture subtle hesitations, even when the boxes are ultimately opened.

Although our overall cohort is larger than the average sample size used in judgement bias tasks conducted in primates (on average less than ten individuals tested; see Machado et al., 2026 for a recapitulative table), we tested a limited number of individuals per species, especially in capuchins. From a statistical standpoint, limited power is common in studies on judgement bias, and even large effect sizes may sometimes be only marginal yet still interpreted as informative results (Winkler et al., 2025). In previous studies on capuchins, judgement bias effects have been reported only on long-term welfare indicators such as overall rates of grooming or stereotypic behaviours (Pomerantz et al., 2012; Schino et al., 2016). For example, Schino and collaborators (2016) reported that cumulative grooming received rates, but not immediate differences in grooming received, predict judgement bias. If proximal effects are not reflected in capuchins’ responses to ambiguity, this could explain why our short-term manipulation appears not significantly effective. Studies combining short and long terms effects, as previously conducted on pigs (Hintze et al., 2025), would thus help build a more comprehensive understanding of judgment bias.

### Some limitations and practical recommendations

#### Effect of judgement bias at the individual level

Although our experimental protocol has proven its effectiveness based on the significant results we observed at the group level for macaques, it has certain limitations regarding the conclusions we can draw at the individual level. Regarding the probability of opening the near negative probe, only seven out of 11 macaques and four out of eight capuchins responded as predicted by the induced affective states. Group-level studies benefit from aggregating data, but actual welfare assessments require individual-level reliability, as each animal assessed must be able to be evaluated on their own. Simple, easily detectable individual differences, rather than complex statistical models, may better suit practical implementation. In that regard, latencies proved more encouraging, as all but three macaques and all but one capuchin exhibited a pattern consistent with the affective state induced. Nevertheless, the inability to confirm that individual results reflect short-term states represents one of the main limitations of this study, as it currently limits its practical application. Individual effects might also be harder to interpret than group level effects, as stable factors such as personality traits, susceptible to impact animal cognition (Carere and Locurto, 2011; Padrell et al., 2020), could bias individuals’ response. Indeed, although still untested in primates, research conducted on other species has shown personality traits such as fearfulness or anxiety to be linked to pessimistic responses in judgment bias tasks (Barnard et al., 2018; Lecorps et al., 2018). Future studies, especially those conducted on small sample size, could benefit from taking such traits into account.

### Training and testing design

Across sessions, we reported a heterogeneous decline in the opening rate of boxes between conditions. To maintain ambiguity and comparability, we thus recommend limiting the presentation of probes (below about 20 presentations, consistent with our results and prior studies; Machado et al., 2026; Schino et al., 2016). As we observed a higher rate of interaction with the first presented cue than the others, we also recommend beginning sessions with a conditioned stimulus to neutralize novelty effects. Our results showed that the unrewarded boxes (negative and probes) were opened less and less across repeated presentation within a session, we thus also advise limiting the number of presentations of ambiguous probe per session.

One limitation of our training procedure is that the negatively conditioned stimulus retained some residual ambiguity throughout the tests. This problem could potentially be resolved by pairing this stimulus with a mildly negative outcome rather than a neutral one (such as a puff of air, e.g. Jones et al., 2018), but this is a welfare trade-off we were unwilling to make and that might also cause participants to lose motivation. For the same reasons, we used only stimuli that caused mild discomfort for the negative condition, which may have limited the intensity of the effect. Future studies should validate these induced states from a behavioural perspective (Adriaense et al., 2021), as a more pronounced manipulation might have produced stronger responses. Conducting opportunistic tests in negatively valenced situations, such as after necessary husbandry procedures (Bethell et al., 2012) or external unexpected conditions (for example a bird of prey flying over the enclosure), could also be a promising avenue of research (although limited control over the situation might restrict the accuracy of these measurements).

### Conclusions

This study proposed an accessible protocol that allowed to successfully train two cohorts of different species of primates and required only a moderate time investment for assessing judgement biases. The effect of induced affective states on judgement biases were observed in macaques but not in capuchins. As is often the case in judgement bias tasks, the progression of sessions attenuated the detectable effect, and we recommend keeping the number of sessions using the same probe to an absolute minimum. We believe that research on affective states should continue to focus on accessible judgement bias tasks such as this one, refining protocols with the aim of practical application relevant to animal welfare.

## Supporting information

Supplementary Additional considerations

Graphical abstract

## Acknowledgments

The authors are grateful to the Simian Laboratory Europe (SILABE, University of Strasbourg, France) for supporting this research and hosting their team. They warmly thank the animal care and welfare staff for their valuable assistance throughout the project, as well as the technical team for their help in designing the device. They also sincerely thank the interns for their dedication and significant contributions to this study: Zazie Benoit-Delaby, Sarah Piaugeard, Taya Chanou and Mathilde Haeuser for their help with the experimental procedures, with the latter two also having participated in the video coding. A special thank goes to Manon Serda for creating the illustrations present in this article.

## Declaration of generative AI and AI-assisted technologies in the manuscript preparation process

During the preparation of this work, the authors used the AI tool Claude (Anthropic) to assist with R code development and debugging. The authors reviewed and edited the content as needed and take full responsibility for the content of the published article.

## CRediT authorship contribution statement

**Aurore Serda:** Conceptualization, Methodology, Formal analysis, Investigation, Writing - Original Draft, Visualization. **Ruta Vaicekauskaite**: Conceptualization, Methodology, Writing - Review & Editing. **Charlotte Canteloup:** Methodology, Writing - Review & Editing, Supervision. **Hélène Meunier**: Methodology, Writing - Review & Editing, Supervision.

## Notes

### Competing Interest Statement

The authors have declared no competing interest.

