## Supplementary Table S1 for "Developing an accessible judgement bias task for non-human primates"

### Supplementary Table S1: Summary of the individuals

*Summary of the individuals included in the study: age, sex, species, group composition, group size, and housing conditions.*

| Subject | Age (years) | Sex (M/F) | Species | Group Type | Group Size | Housing |
|---|---|---|---|---|---|---|
| Walt | 17 | M | *Macaca tonkeana* | All-male | 6 | Natural outdoor enclosure, ~1364m$^2$ outdoor + 10m$^2$ indoor access |
| Yang | 15 | M | | | | |
| Eric | 7 | M | | All-male | 3 | Concrete outdoor enclosure, ~16m$^2$ outdoor + 14m$^2$ indoor access |
| Barnabé | 10 | M | | | | |
| Abricot | 11 | M | | | | |
| Alaryc | 12 | M | | All-male | 2 | Concrete outdoor enclosure, ~16m$^2$ outdoor + 10m$^2$ indoor access |
| César | 9 | M | | | | |
| Yannick | 16 | M | | Mixed | 20 | Natural outdoor enclosure, ~3788m$^2$ outdoor + 20m$^2$ indoor access |
| Indigo | 4 | M | | | | |
| Iron | 3 | M | | | | |
| Nereis | 26 | F | | | | |
| Franklin | 14 | M | *Sapajus apella* | All-male | 3 | Concrete outdoor enclosure, ~28m$^2$ outdoor + 7.5m$^2$ indoor access |
| Kiri | 12 | M | | | | |
| Brugnon | 10 | M | | | | |
| Popeye | 23 | M | | All-male | 3 | Concrete outdoor enclosure, ~13.5m$^2$ outdoor + 4.2m$^2$ indoor access |
| Conan | 9 | M | | | | |
| Balin | 10 | M | | | | |
| Dorianne | 8 | F | | All-female | 2 | Natural outdoor enclosure, ~9m$^2$ outdoor + 8m$^2$ indoor access |
| Kaina | 12 | F | | | | |
