## Supplementary Methods S1 for "Developing an accessible judgement bias task for non-human primates"

### Supplementary Methods S1: Colorimetric quantification of cue position

The colours of the cues were chosen to be as equidistant as possible from the stimuli at each extremity. To verify the expected order and approximate spacing of the two ambiguous probes, the perceptual colour distance between each probe and the two trained cues (CS-, white; CS+, black) was quantified using the CIEDE2000 colour-difference formula (Sharma et al., 2005), via the online calculator at ColorMine.org (https://colormine.org/delta-e-calculator/cie2000). The colour of the cues were as follows: CS- (white) = rgb(233, 224, 210), P- (light grey) = rgb(144, 148, 149), P+ (dark grey) = rgb(87, 92, 95), CS+ (black) = rgb(10, 10, 10).

For each probe, the ΔE was calculated relative to the white and black reference colours, then normalised on a scale from 0 (white) to 1 (black):

$$Position = \Delta E(white) / [\Delta E(white) + \Delta E(black)]$$

| **Cue** | **ΔE to white** | **ΔE to black** | **Position (0-1)** |
|---|---|---|---|
| P- | 22.32 | 46.16 | 0.33 |
| P+ | 47.75 | 24.10 | 0.66 |

To make interpretation easier, cue positions have been rounded off further, the final cue positions are as follows: CS− = 0, P− = 0.3, P+ = 0.7, CS+ = 1.
