## Supplemental Data 1 for "Developing an accessible judgement bias task for non-human primates"

### Supplementary: Statistical results

#### Supplementary Table S2: Sensitivity analyses.

*(a) Macaques: subjects tested in both experimental phases (n = 5) compared with the full sample (n = 11). (b) Capuchins: subjects excluding two individuals who exceeded the CS- opening threshold (n = 6) compared with the full sample (n = 8). Values shown are Type II Wald $\chi^2$ (binomial models) or Type II likelihood-ratio $\chi^2$ (Cox models) and p values for the key term(s) of each model.*

##### (a) *Macaca tonkeana* (n = 5 vs. full sample, n = 11)

| Model | Term | $\chi^2$ (full, n=11) | p (full) | $\chi^2$ (n=5) | p (n=5) |
|---|---|---|---|---|---|
| Discrimination | cue | 242.45 | **<0.001** | 114.42 | **<0.001** |
| | session number | 1.44 | 0.230 | 3.75 | 0.053 |
| Discrimination (latency) | cue | 654.02 | **<0.001** | 395.25 | **<0.001** |
| | cue×session | 23.12 | **<0.001** | 11.49 | **0.009** |

##### (b) *Sapajus apella* (n = 6 vs. full sample, n = 8)

| Model | Term | $\chi^2$ (full, n=8) | p (full) | $\chi^2$ (n=6) | p (n=6) |
|---|---|---|---|---|---|
| Discrimination | cue | 226.27 | **<0.001** | 184.17 | **<0.001** |
| Discrimination (latency) | cue×session | 28.61 | **<0.001** | 20.83 | **<0.001** |
| Affective state, phase L | condition | 0.16 | 0.689 | 0.02 | 0.880 |
| Affective state, phase D | condition | 1.50 | 0.220 | 2.25 | 0.133 |
| Affective state, phase L (latency) | condition | 2.89 | 0.089 | 0.65 | 0.421 |
| Affective state, phase D (latency) | condition | 0.03 | 0.854 | 0.02 | 0.885 |

#### Supplementary Table S3: Estimated marginal means by cue and species.

*Post-hoc estimated marginal means by cue and species for the binomial GLMM on box-opening probability (GLMM1_M, GLMM1_S; at first_trial = 0).*

| Species | Cue | Probability (95% CI) |
|---|---|---|
| *M. tonkeana* | CS− | 0.062 (0.031–0.119) |
| | P− | 0.198 (0.111–0.329) |
| | P+ | 0.448 (0.268–0.642) |
| | CS+ | 0.988 (0.970–0.995) |
| *S. apella* | CS− | 0.098 (0.045–0.201) |
| | P− | 0.407 (0.227–0.616) |
| | P+ | 0.479 (0.283–0.682) |
| | CS+ | 0.994 (0.982–0.998) |

#### Supplementary Table S4: Remaining pairwise contrasts of latency to open

*Pairwise contrasts of latency to open of non-consecutive comparisons along the ambiguity gradient, by session and species, from the mixed-effects Cox model (Cox1_M, Cox1_S; Holm-adjusted for 6 comparisons per session). Each cell reports ratio / SE / p values. Ratios above 1 indicate faster opening for the first-listed cue.*

| Species | Contrast | Session 1 | Session 2 | Session 3 | Session 4 | Session 5 |
|---|---|---|---|---|---|---|
| M. tonkeana | CS+ / CS− | 9.18 / 1.63 / **<0.001** | 13.34 / 1.73 / **<0.001** | 19.38 / 2.51 / **<0.001** | 28.17 / 4.97 / **<0.001** | 40.94 / 10.00 / **<0.001** |
| | CS+ / P− | 5.67 / 1.02 / **<0.001** | 7.00 / 0.91 / **<0.001** | 8.63 / 1.01 / **<0.001** | 10.64 / 1.60 / **<0.001** | 13.13 / 2.74 / **<0.001** |
| | P+ / CS− | 2.71 / 0.73 / **<0.001** | 3.29 / 0.60 / **<0.001** | 4.00 / 0.69 / **<0.001** | 4.87 / 1.19 / **<0.001** | 5.92 / 2.08 / **<0.001** |
| S. apella | CS+ / CS− | 10.15 / 1.69 / **<0.001** | 12.04 / 1.46 / **<0.001** | 14.29 / 1.54 / **<0.001** | 16.94 / 2.30 / **<0.001** | 20.10 / 3.76 / **<0.001** |
| | CS+ / P− | 3.14 / 0.54 / **<0.001** | 3.83 / 0.47 / **<0.001** | 4.68 / 0.50 / **<0.001** | 5.71 / 0.75 / **<0.001** | 6.96 / 1.27 / **<0.001** |
| | P+ / CS− | 4.11 / 0.81 / **<0.001** | 3.33 / 0.47 / **<0.001** | 2.70 / 0.35 / **<0.001** | 2.19 / 0.38 / **<0.001** | 1.77 / 0.43 / **0.036** |

#### Supplementary Table S5: Individual results

*Individual mean box-opening probability and latency to open (seconds) for P- trials, by condition and subject (n = 19). Δ indicates the change from negative to positive condition (percentage points for opening probability; seconds for latency); green indicates change in the expected direction (higher opening probability / shorter latency under positive condition), red indicates the opposite.*

| | Opening probability | | Latency (s) | |
|---|---|---|---|---|
| **Subject** | **Negative** | **Positive (Δ)** | **Negative** | **Positive (Δ)** |
| ***Macaca tonkeana*** | | | | |
| Abricot | 0.58 | 0.60 (+2 pp) | 2.68 | 2.35 (-0.33 s) |
| Alaryc | 0.07 | 0.20 (+13 pp) | 4.40 | 3.80 (-0.60 s) |
| Barnabé | 0.13 | 0.40 (+27 pp) | 4.69 | 3.41 (-1.28 s) |
| César | 0.27 | 0.13 (-13 pp) | 3.89 | 3.90 (+0.01 s) |
| Eric | 0.75 | 0.71 (-4 pp) | 1.84 | 1.62 (-0.22 s) |
| Indigo | 0.53 | 0.87 (+33 pp) | 2.47 | 2.06 (-0.41 s) |
| Iron | 0.20 | 0.40 (+20 pp) | 4.17 | 3.92 (-0.25 s) |
| Nereis | 0.08 | 0.17 (+8 pp) | 2.67 | 3.84 (+1.17 s) |
| Walt | 0.21 | 0.00 (-21 pp) | 3.59 | 4.67 (+1.08 s) |
| Yang | 0.13 | 0.07 (-7 pp) | 4.38 | 4.34 (-0.05 s) |
| Yannick | 0.07 | 0.27 (+20 pp) | 4.68 | 3.80 (-0.88 s) |
| ***Sapajus apella*** | | | | |
| Balin | 0.53 | 0.53 (+0 pp) | 2.30 | 1.30 (-1.00 s) |
| Brugnon | 0.27 | 0.53 (+27 pp) | 3.72 | 1.71 (-2.02 s) |
| Conan | 0.40 | 0.13 (-27 pp) | 3.25 | 3.01 (-0.24 s) |
| Dorianne | 0.07 | 0.33 (+26 pp) | 3.91 | 2.80 (-1.11 s) |
| Franklin | 0.87 | 0.87 (+0 pp) | 1.32 | 0.69 (-0.62 s) |
| Kaina | 0.40 | 0.47 (+7 pp) | 3.05 | 2.23 (-0.82 s) |
| Kiri | 0.47 | 0.67 (+20 pp) | 2.62 | 1.98 (-0.64 s) |
| Popeye | 0.73 | 0.60 (-13 pp) | 2.36 | 2.98 (+0.62 s) |

#### Supplementary Table S6: Successive trial presentation effects

*Model-predicted box-opening probability (95% CI) across successive within-session presentations of the same cue. (a) Macaques: by cue, given a significant cue×presentation-number interaction (GLMM3_M: $\chi^2_3 = 10.03$, $p = 0.018$). (b) Capuchins: averaged across cues, as this interaction was not significant (GLMM3_S: $\chi^2_3 = 7.44$, $p = 0.059$).*

##### (a) *Macaca tonkeana*

| Cue | Presentation 1 | Presentation 2 | Presentation 3 |
|---|---|---|---|
| CS− | 0.147 (0.069–0.287) | 0.050 (0.023–0.106) | 0.016 (0.006–0.043) |
| P− | 0.389 (0.219–0.591) | 0.179 (0.092–0.318) | 0.069 (0.029–0.158) |
| P+ | 0.582 (0.333–0.795) | 0.472 (0.273–0.680) | 0.364 (0.173–0.611) |
| CS+ | 0.990 (0.962–0.997) | 0.990 (0.975–0.996) | 0.991 (0.971–0.997) |

##### (b) *Sapajus apella*

| | Presentation 1 | Presentation 2 | Presentation 3 |
|---|---|---|---|
| All cues (averaged) | 0.759 (0.570–0.883) | 0.663 (0.463–0.818) | 0.552 (0.345–0.742) |
