## Supplementary Figure S1 for "Developing an accessible judgement bias task for non-human primates"

**Supplementary Figure S1: Schematic decision tree for trial flow during the box discrimination training and tests.**

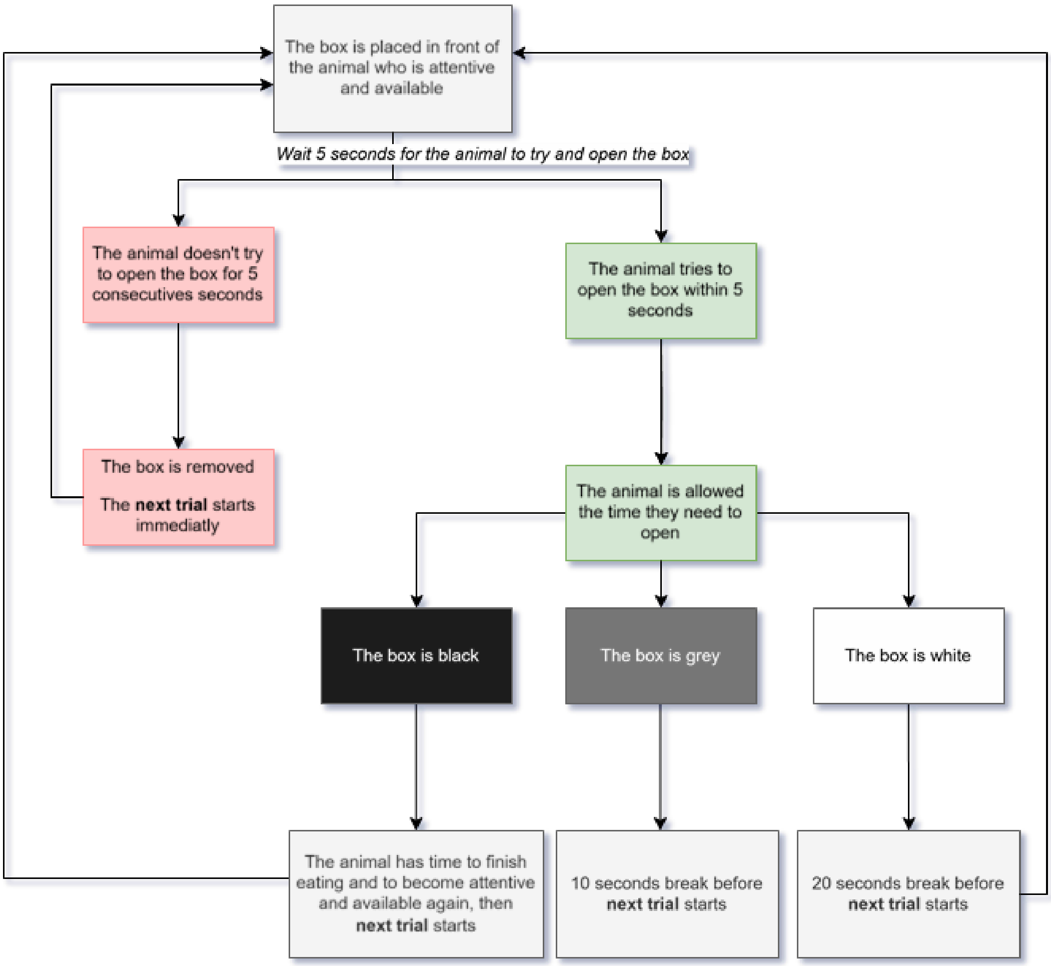
