## Supplementary Additional considerations for "Developing an accessible judgement bias task for non-human primates"

### Additional considerations

#### Effect of session on latency to open the box according to cue

The effect of session on the latency to open boxes differed between CS+ boxes and CS- boxes as well as between CS+ boxes and P- boxes in macaques (CS+ vs CS-: estimate = 0.37, $p < 0.001$; CS+ vs P-: estimate = 0.21, $p = 0.035$), whereas CS- boxes, P- boxes, and P+ boxes were not impacted differently from one another (all $p > 0.3$). In capuchins, the progression of sessions impacted the latency to open P+ boxes significantly differently from CS+ boxes ($z = 2.53$, $p < 0.001$) and from P- boxes ($z = 2.35$, $p = 0.025$), while CS- boxes, P- boxes, and CS+ boxes' latencies did not evolve with sessions differently from one another (all $p > 0.07$).

#### On the significant effect of condition on overall latency for macaques

In contrast to the effect on the box opening rate, only a general effect (cue-averaged) of the condition on latency was found for the first phase in macaques, with faster openings under positive condition. This discrepancy could be explained by the latency close to the ceiling, and with low-variance observed for the positive conditioned stimulus, which weakens the statistical power required to detect cue-specific patterns. By comparison, latency measures may also be more sensitive to general disturbance or motivation than binary responses (Bateson and Nettle, 2015), reflecting motivational states rather than a bias. However, a comparable study on marmosets (*Callithrix jacchus*) showed that both measures reflected bias in a similar manner (Gordon and Rogers, 2015).
