## Supplementary figures and images for "Developing an accessible judgement bias task for non-human primates"

### Graphical abstract

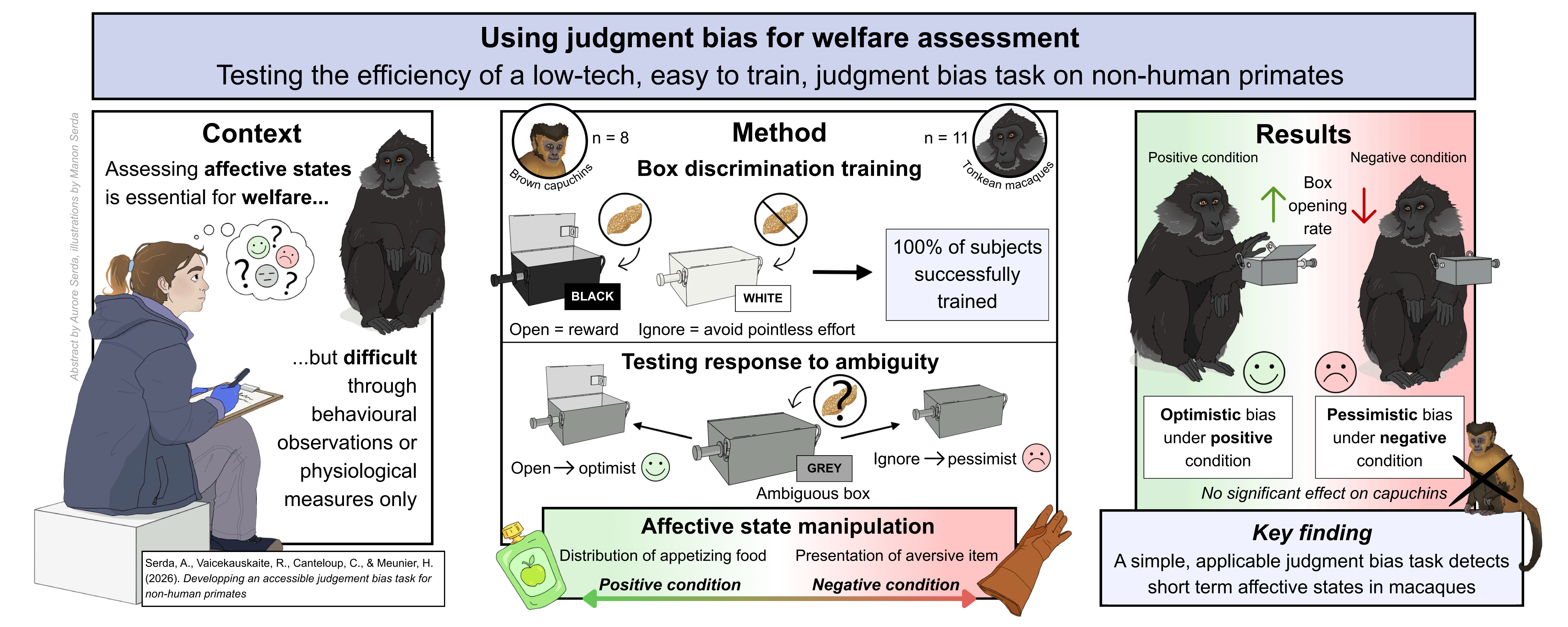
